# Plasma concentrations of progesterone in the preceding estrous cycle are associated with the intensity of estrus and fertility of Holstein cows

**DOI:** 10.1101/2021.03.01.433335

**Authors:** A.M.L. Madureira, T.A. Burnett, S. Borchardt, W. Heuwieser, C.F. Baes, J.L.M. Vasconcelos, R.L.A. Cerri

## Abstract

The aim of this study was to determine the association between concentrations of progesterone **(P4)** during previous the estrous cycle with the intensity of spontaneous or estrogen-induced estrous expression and pregnancy per artificial insemination **(P/AI)**. A total of 1,953 AI events from lactating Holstein cows were used, consisting of 1,289 timed AI events from experiment 1 **(Exp. 1)** and 664 AI events from experiment 2 **(Exp. 2)**. In Exp. 1, cows were bred after a timed AI protocol based on estradiol and P4. In Exp. 2 animals were bred upon spontaneous estrus detection. In both experiments cows were continuously monitored by an automated activity monitor **(AAM)**. The intensity of estrous expression was determined for each event and classified as either high or low intensity using the median of each experiment. Blood samples were collected for P4 analysis in Exp. 1 at –4 d, 0 d, and 7 d relative to timed AI, and in Exp. 2 immediately following estrus (0 d), 7 d, 14 d, and 21 d post-AI. Concentration of P4 was classified as greater or lower according to the median value in each experiment. Cows with lower concentrations of P4 at AI had greater estrous expression in Exp. 1 (363.6 ± 5.2 vs. 275.9 ± 8.0 % relative increase) and Exp. 2 (76.7 ± 1.9 vs. 67.4 ± 4.7 index; and 12.5 ± 0.5 vs. 9.3 ± 1.8 h). Cows with a greater intensity of estrous expression at timed AI had greater concentrations of P4 at –4 d than cows with lower intensity estrus or no estrous expression (4.6 ± 0.2 vs. 3.6 ± 0.2 vs. 3.7 ± 0.2 ng/mL). Cows with greater concentrations of P4 at –4 d had greater P/AI (32.8 ± 4.4 vs. 22.4 ± 4.5 %), whereas cows with lesser concentrations of P4 at d0 for either timed AI (35.2 ± 3.4 vs. 19.6 ± 4.6 %) or spontaneous estrus (31.8 ± 2.8 vs. 23.4 ± 3.2 %) had greater P/AI. Cows with greater concentrations of P4 7 d post-timed AI had greater P/AI compared with cows that had lower concentration of P4 (39.1 ± 2.9 vs. 24.7 ± 2.6 %). Similarly, cows that had lower concentrations of P4 at 7 d, 14 d and 21 d post-spontaneous estrus tended to have lower P/AI when compared with cows with greater concentrations of P4. Overall, concentrations of P4 prior to and at AI were associated with greater estrous intensity and P/AI at both spontaneous and timed AI events.

## Introduction

Visual estrus detection is difficult due to the shorter duration and lower intensity of estrous expression [1–3] observed in modern dairy cows. However, automated activity monitors **(AAM)** have proven to be efficient and reliable technologies as an alternative estrus detection strategy over visual detection [4].

In a series of studies using different AAM systems, pregnancy per artificial insemination (**P/AI**) was reported to be greater for cows with more intense estrus [5–7]. Pereira et al. [8] reported that cows expressing estrus at the end of a timed AI protocol based on estradiol **(E2)** and progesterone **(P4)** had improved fertility using timed AI (38.9% vs. 25.5%) and embryo transfer (46.7% vs. 32.7%) compared with those that did not express estrus. Furthermore, pregnancy loss was lower for cows that expressed estrus at timed AI (14.4% vs. 20.1%) and prior to embryo transfer (18.6% vs. 22.7%; [8]).

Pereira et al. [8] reported that cows that did not express estrus at the end of a timed AI protocol were more likely to have had higher concentrations of P4 at AI compared with cows that expressed estrus. This could be associated with incomplete luteal regression during the proestrus period has often been reported [9–11]. In these studies, cows with slightly higher plasma concentrations of P4 at AI, possibly caused by the absence of complete corpus luteum (**CL**) regression, had decreased fertility. It is therefore possible that an elevated P4 concentration directly at AI is detrimental, while elevated P4 levels at other timepoints before and after AI are beneficial. For example, Madureira et al. [12] noted that the concentration of P4 at 7 d post-AI was greater in cows with intensive estrous expression, which is correlated with higher P/AI.

Numerous studies have demonstrated the effects of P4 on fertility at and immediately after AI. However, literature regarding the association of the sequence of circulating P4 before AI, at AI, and post-AI with the intensity of estrous expression in cattle is limited. The overall objective of this study was to determine the association between the concentrations of P4 at different phases of the preceding estrous cycle and the intensity of spontaneous or estrogen-induced estrous expression (detected by AAM), and on P/AI. We hypothesized that higher concentrations of P4 at different phases of diestrus, coupled with lower concentrations of P4 at AI, would be associated with greater estrous expression and improved P/AI.

## Material and methods

Two experiments were conducted at two different locations. Experiment 1 (**Exp. 1**) was conducted on a commercial farm in the state of São Paulo, Brazil (latitude: 22°21’25” S; longitude: 47°23’03” W) while Experiment 2 (**Exp. 2**) was conducted at the University of British Columbia’s Dairy Education and Research Centre in Agassiz, Canada (49°13’59” N, 121°46’01” W). In Exp. 1, estrus detection was monitored by an AAM based on steps/h, and cows were bred after a synchronization protocol based on E2 and P4. In Exp. 2, estrus detection was monitored by an AAM based on an activity index, and all cows were artificially inseminated after spontaneous estrus. The objective of Exp. 1 was to determine the association between the concentrations of P4 at different phases of the estrous cycle, the intensity of estrogen-induced estrous expression, and on P/AI. The objective of Exp. 2 was to confirm the findings of Exp.1 with AI based on naturally occurring estrus and to elucidate the relationships of the hormonal profile after AI with fertility also considering size of the follicle. The University of British Columbia’s Animal Care protocol related with the current study was A18-0315 (valid for both experiments). The practices outlined in the Guide for the Care and Use of Agricultural Animals in Agricultural Research and Teaching [13], and the Canadian Council on Animal Care [14] were used for the approval of all experimental procedures as part of the local requirements.

### Animals, housing and management

Cows from Exp. 1 were enrolled at the beginning of a timed AI protocol (Figure 1A) from a commercial herd with an average 305-d mature equivalent milk yield of 11,438 kg/cow and approximately 1,700 lactating Holstein dairy cows. Cows from Exp. 2 were enrolled upon the detection of spontaneous estrus (Figure 1B) from an experimental herd with an average 305-d mature equivalent milk yield of 12,799 kg/cow and approximately 270 lactating Holstein dairy cows. At the time of enrollment, body condition score **(BCS)** was assessed (1 to 5 scale at 0.25 increments; [15]) in both experiments. Milk production was measured daily at each milking (AfiLite, Kibbutz, Afikim, Israel) in Exp. 1 between –11 d and 0 d (day of timed AI) of the experiment and the average daily milk production during this period was used for analysis. In Exp. 2, milk production was recorded daily and averaged from 2 days before until 2 days after the detection of estrus.

**Figure I.**
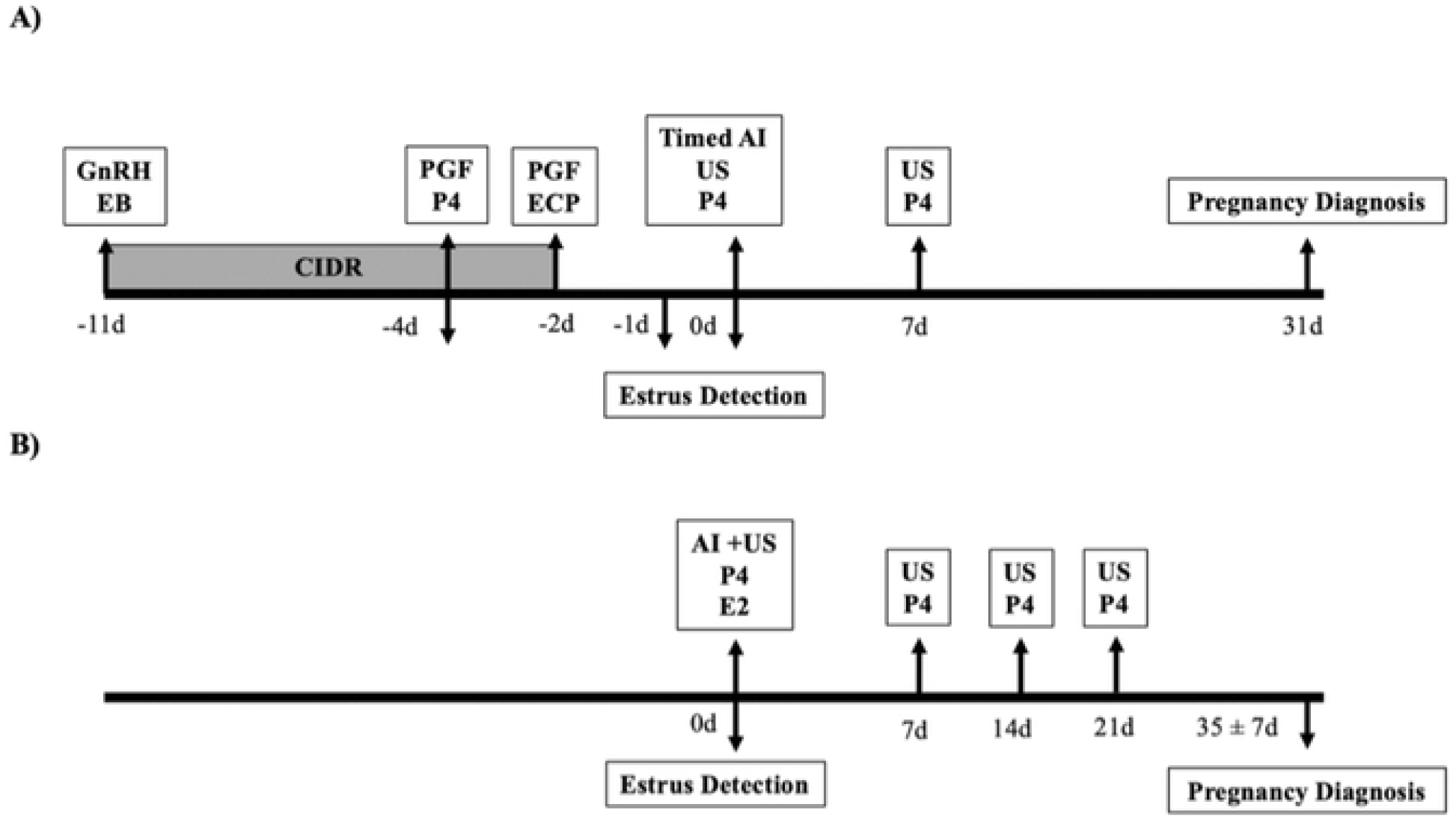
Schematic of the experimental design for Experiment I (Exp. I; panel A) and Experiment 2 (Exp.2; panel B). EB (estradiol benzoate; 2 mg, Gonadiol, Zoetis, Sao Paulo, Brazil), GnRH (gonadorelin diacetate; 100 11g, Cystorelin, Merial, Sao Paulo, Brazil), CIDR (intravaginal progesterone i1nplant; 1.9 g progesterone; CIDR, Zoetis, Sao Paulo, Brazil), PGF (dinoprost tromethamine; 25 mg, Lutalyse, Zoetis, Sao Paulo, Brazil), ECP (estradiol cypionate; I mg, E.C.P., Zoetis, Sao Paulo, Brazil), Timed Al (artificial insen1ination), Al (artificial insemination, followed by spontaneous estrus using the a111/pmntle; Exp. 2 only), US (examination of ovaries with ultrasonography), P4 (collection of blood sample for analysis of progesterone concentration), E2 (collection of blood sample of analysis of estradiol concentrations). Automated detection of estrus was done using a leg-mounted pedometer in Exp. 1 (Afimilk Pedometer Plus Tags; Afimilk, Kibbutz Afikim, Israel) and a neck-mounted accelero1neter in Exp. 2 (Heatime®, SCR Engineers, Israel).

In Exp. 1 cows were housed in a cross-ventilated free-stall barn, while cows in Exp. 2 were housed in a naturally ventilated free-stall barn. Milking was performed three times daily (at approximately 0500, 1300, and 2100 h) in Exp. 1, while milking was performed twice daily (at approximately 0500 and 1500 h) in Exp. 2. Cows were fed a total mixed ration **(TMR)** three times daily in Exp. 1 and twice daily in Exp. 2. The TMR was formulated to meet or exceed the requirements of a lactating Holstein cow producing 40 kg/day of 3.5% fat corrected milk [16] and pushed up three times daily on both locations. Water and TMR were available for *ad libitum* intake.

### Automated activity monitors

Cows were continuously monitored by a leg-mounted pedometer (AfiPedometer Plus Tag; AfiMilk^®^, Israel) in Exp. 1. The pedometer was attached to the right back leg of each cow on the day of calving where it remained throughout the entire experimental period. The data from the AAM were used to calculate the relative increase in physical activity. To determine the relative increase the following formula was used: [(steps/h at estrus - steps/h at baseline)/steps/h at baseline] * 100]. Maximum steps/h at estrus was considered as the greatest steps/h count of the entire estrus event and the baseline was defined as the average steps/h for the 7d prior to AI. Cows that exceeded a relative increase of 100 % on either –1 d or 0 d relative to timed AI were considered to be in estrus at the time of AI, whereas those with a relative increase < 100 % were classified as not having an alert from the AAM, as described by Madureira et al. [12].

In Exp. 2 a neck-mounted accelerometer (Heatime^®^, SCR Engineers, Netanya, Israel) was placed on the upper left side at the cranial portion of the cow’s neck using a collar on the day of calving and it was removed if a positive pregnancy diagnosis was detected. The activity data from each cow were computed into an index value that ranged from 0 to 100 through the system’s proprietary algorithm and a threshold of 35 was used as an alert for an estrus event. The maximum increase in physical activity index and the interval from the onset to the end of the alert (h above threshold) was calculated using an Excel macro (Excel; Microsoft Corporation, Redmond, WA) to identify estrus events from the exported backup files.

In order to simplify the description of estrus intensity categories, the median value of activity value from the two AAM systems in Exp. 1 and Exp. 2, as well as the total duration of the estrus episodes in Exp. 2 were used; all three were later classified using the median. Estrus events were classified as High Intensity and Low Intensity using the threshold of 300 % relative increase in Exp. 1, and 80.5 Index in Exp. 2. The duration of estrus was classified as Long and Short using the threshold of 8 h.

### Synchronization protocol, artificial insemination, ultrasonography, and pregnancy diagnosis

#### Experiment 1

All cows were enrolled onto a synchronization protocol based on progesterone and estradiol as described by Pereira et al. [17] and detailed in Figure 1. Timed AI was performed using commercially frozen–thawed semen by two trained technicians. Ovaries of a subset of cows were examined by ultrasonography (Honda Electronics, Toyohashi, Japan) using a 7.5 MHz linear-array rectal transducer on the day of AI (0 d; assessment of two largest follicles; n = 541) and 7 days post-AI (7 d; presence or absence of a CL to confirm ovulation; n = 741). Pregnancy diagnosis was performed using ultrasonography on 31 d ± 3 d after timed AI for all cows. A cow was considered pregnant if an embryo with heartbeat was present. Pregnancy per AI was calculated by dividing the number of cows that were pregnant 31 d after AI by the number of animals inseminated after the timed AI protocol.

#### Experiment 2

Cows were enrolled onto the study when they were identified as in estrus by the neck mounted automated activity monitor. All cows had their ovaries examined by ultrasonography (Aloka SSD-500, Aloka Co. Ltd, Wallingford, CT, USA) using a 7.5 MHz linear rectal transducer at the time of an estrus alert as well as at 7 d, 14 d and 21 d post-AI. Presence and diameter of a CL and/or diameter of the largest follicles were measured and recorded at each assessment. Cows were artificially inseminated upon estrus using the a.m./p.m. rule. True events from the AAM alerts were considered based on the presence of a pre-ovulatory follicle larger than 15 mm in diameter and absence of an active CL (greater than 25 mm in diameter). All cows were examined with an ultrasound at 35 ± 7 d post-AI for pregnancy diagnosis, cows that had an embryo with a heartbeat were considered positive for pregnancy.

### Blood sampling, analyses of estradiol and progesterone concentrations

#### Experiment 1

Blood samples were harvested from the median coccygeal vein or artery into commercial blood collection tubes (BD Vacutainer Serum Tubes, 10 mL; Becton Dickinson, Franklin Lakes, NJ). Blood samples were collected on -4 d (n = 312), 0 d (n = 923) and 7 d (n = 780) relative to timed AI. All samples were immediately placed on ice and then centrifuged at 3,000 × g at 4°C for 30 min for serum collection and stored at -20°C until analyzed for P4 concentration. Serum P4 concentrations were analyzed using a chemiluminescent enzyme immunoassay (Immulite 1000; Siemens Medical Solutions Diagnostics, Los Angeles, CA) that was previously validated [18,19]. The intra- and inter-assay coefficients of variation **(CV)** were 5.1 and 5.2 %, respectively. The minimum detectable concentration was 0.1 ng/mL of P4.

#### Experiment 2

Blood samples were harvested from the median coccygeal vein or artery utilizing Vacutainer tubes (10 mL; Becton & Dickinson Vacutainer systems, Rutherford, NJ) with K_2_EDTA. Blood samples were collected immediately following a detected estrus alert (0 d; n = 553) by the AAM system and at 7 d (n = 142), 14 d (n = 148) and 21 d (n = 138) post-AI for P4 analysis. Only samples collected at the estrus alert (0 d; n = 617) were analyzed for E2. All samples were immediately placed on ice, transported to the laboratory and centrifuged at 2,700 x g for 15 min for separation of plasma. Plasma samples were then stored at -80 °C to be analyzed for E2 and P4 concentrations. Plasma concentrations of E2 were determined using a double-antibody I125-based assay developed as described by Burke et al. [20] with modifications. Average intra-assay CV was 4.2 %, and inter-assay CV for pooled plasma samples containing 2.7, 5.8, and 12.3 pg/mL of E2 were 6.3 %, 5.2 % and 4.1 %, respectively. The average sensitivity of the E2 assay was 1.1 pg/mL, as described by Madureira et al. [21]. Plasma P4 concentrations were measured using a commercial ELISA kit (Ovucheck Plasma; Biovet, St-Hyacinthe, Quebec; Broes and LeBlanc, [22]. The range of quantification of the test is 0.55 – 10.45 ng/mL, and the intra-assay CV in the present study was 10.8 %.

### Statistical analysis

In Exp. 1, the sample size (n = 281 per group) was calculated to identify a difference of 15% (20% vs. 35%) in pregnancy per AI, between cows expressing high and low intensity of estrus at AI with 95% confidence and 80% power. This sample size was also sufficient (sample size = 83 per group) to identify a 25 % increase in activity (300 vs 375%), expecting a SD of 130 steps/h between cows with different P4 concentrations prior to AI (-4 d) and at the day of AI (0 d;); this sample size was also sufficient to identify a difference of 1.0 ng/mL in P4 concentration at 7 d post-AI, expecting a SD of 2.0 ng/mL between cows expressing high and low intensity of estrus at AI (sample size = 63 per group). In Exp. 2, the sample size (n = 80 per group) was calculated to identify a difference of 11 index points, expecting a SD of 18.9 index points for cows with different P4 concentrations at estrus with 95% confidence and 80% power. The sample size (n = 109 per group) was calculated to identify a difference of 1.0 ng/mL in P4 concentration at 7 d, 14 d and 21 d post-AI, expecting a SD of 2.0 ng/mL for cows expressing either high or low intensity of estrus at AI.

Distributions and normality tests were obtained using the Univariate procedure of SAS software, version 9.4 (SAS Institute Inc., Cary, NC). Any variable found not be normally distributed were categorized into class variables. Class variables used for analyses are described below. Parity was divided as cows in first lactation and second lactation or higher (primiparous vs. multiparous), in both experiments. For Exp. 1 and Exp. 2, BCS was categorized as Low (< 2.75), Moderate (2.75 – 3.00), and High (≥ 3.00). Physical activity of estrus episodes in Exp. 1 was categorized by the median maximum activity of estrus events (300 % relative increase) as follows: No Estrus (< 100 % relative increase in activity), Low intensity (100 – 299 % relative increase in activity), and High intensity (≥ 300 % relative increase in activity). A 100 % relative increase was used as the estrus threshold as previously described by Madureira et al. [12]. In Exp.2, the activity at time of alert was categorized relative to the median as High intensity (≥ 80.5 index) and Low intensity (< 80.5 index). Duration of the estrus event was also categorized using the median as Long duration (≥ 8h) and Short duration (< 8h). In Exp. 1, concentrations of P4 were classified as lesser or equal to the median and greater than the median at each sampling time using the following values: -4 d prior to timed AI (3.04 ng/mL); 0 d (at the day of timed AI; 0.1 ng/mL) and at 7 d post-AI (2.96 ng/mL). Concentrations of P4 were categorized by the median in Exp. 2 as having greater or lower concentration as: on 0 d (at the moment of AI; 0.26 ng/mL); on 7 d post-AI (2.65 ng/mL); on 14 d post-AI (3.12 ng/mL); and on 21 d post-AI (6.27 ng/mL). To evaluate the cumulative effect of both the concentration of P4 at -4 d and 0 d on estrous expression and P/AI in Exp. 1, a classification was created according to the median values as described above: **HL** = high concentration of P4 at -4 d and low concentration of P4 at 0 d; **LH =** low concentrations of P4 at -4 d and high concentration of P4 at 0 d; **HH** = high concentration of P4 at -4 d and high concentration of P4 at 0 d; and **LL** = low concentration of P4 at -4 d and low concentration of P4 at 0 d.

Pearson correlations between physiological measurements (e.g., diameter of pre-ovulatory follicle, progesterone concentration, estradiol concentration, DIM, and milk production) and automated activity measurements (e.g., intensity and duration of estrus) were determined. Relative increase in physical activity (Exp. 1), maximum increase in activity (Exp. 2), and total duration were used as continuous dependent variables and were tested for the effects of parity, BCS, milk production, DIM, pre-ovulatory follicle diameter and corpus luteum size by ANOVA using mixed linear regression models (GLIMMIX) in SAS with artificial insemination event as the experimental unit and cow as a random effect. Parity, BCS, milk production, DIM, pre-ovulatory follicle diameter and corpus luteum size were used as independent variables. Diameter of the pre-ovulatory follicle was tested as a continuous dependent variable for the effects of parity, BCS, milk production, relative increase (Exp. 1), activity index (Exp. 2), and duration (Exp. 2), and size of the corpus luteum using mixed linear regression models as described above; separate models were used for each estrous expression measurement. Pregnancy per AI was used as a binomial dependent variable assessed for the same effects listed above with a mixed logistic regression model using the GLIMMIX procedure, where AI event was again used as the experimental unit and cow as the random effect. For the analysis of pregnancy per AI associated with the concentration of P4 at 7 d, 14 d and 21 d post-AI only animals that ovulated were included. Only variables with a *P*-value < 0.15 were kept in the final model. All multivariable models were constructed using the variables as described above and manual backward stepwise elimination was used, where variables were retained with *P* ≤ 0.15. Interactions were tested between all variables selected in the final model and kept if *P* ≤ 0.05. Differences with *P* ≤ 0.05 were considered significant and those between 0.05 > *P* ≤ 0.10 were designated as a tendency and data reported as least squared means where applicable.

## Results

### Animals and number of events

A total of 2,020 AI events from lactating Holstein cows were enrolled. A total of 67 cows had missing estrus intensity data from Exp. 1. A total of 1,289 timed AI events from 984 cows were recorded for Exp.1 with an average milk production of 45.5 ± 10.7 kg/d with approximately 3.5% fat, averaging 117.2 ± 58.2 DIM. In total, 44.2% and 55.8% of cows enrolled were primiparous and multiparous, respectively. In Exp. 2, a total of 664 events from 291 cows were recorded, with an average milk production of 38.4 ± 8.2 kg/d with approximately 4.7% fat, averaging 118.4 ± 52.9 DIM. In Exp. 2 a total of 39.2 % and 60.8% of cows enrolled were primiparous and multiparous, respectively.

### Intensity of estrous expression at AI

The mean relative increase in activity at estrus was 328.7 ± 132.7 % in Exp. 1 and the mean activity index and duration at estrus was 76.6 ± 18.9 index and 12.6 ± 9.7 h, respectively, in Exp. 2. In Exp. 1, 13.7 % (167/1222) of the cows did not display estrus at timed AI (relative increase in activity < 100 %). Parity influenced estrous expression in both experiments as multiparous cows expressed lower relative increase, activity index, and duration at the day of AI (*P* < 0.01; Table 1) when compared with primiparous cows. In Exp. 1, BCS at timed AI tended to be associated with relative increase (*P* = 0.10; Table 1), as cows with lower BCS tended to have lower estrous expression compared with moderate or high BCS. In Exp. 2, BCS was associated with maximum activity index and duration (*P* = 0.01; Table 1), where cows with lower BCS generally had less intense estrous expression. There was no correlation between milk production and relative increase in activity in Exp. 1 (r = - 0.02; *P* = 0.37), but there was a weak negative correlation for both estrous expression measurements (activity index and duration) in Exp. 2 (maximum activity index: r = - 0.20; *P* < 0.01; duration: r = - 0.18; *P* < 0.01).

**Table 1.**
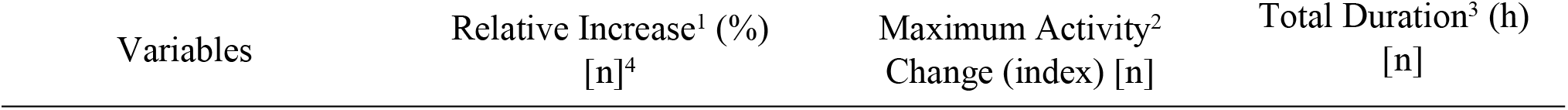
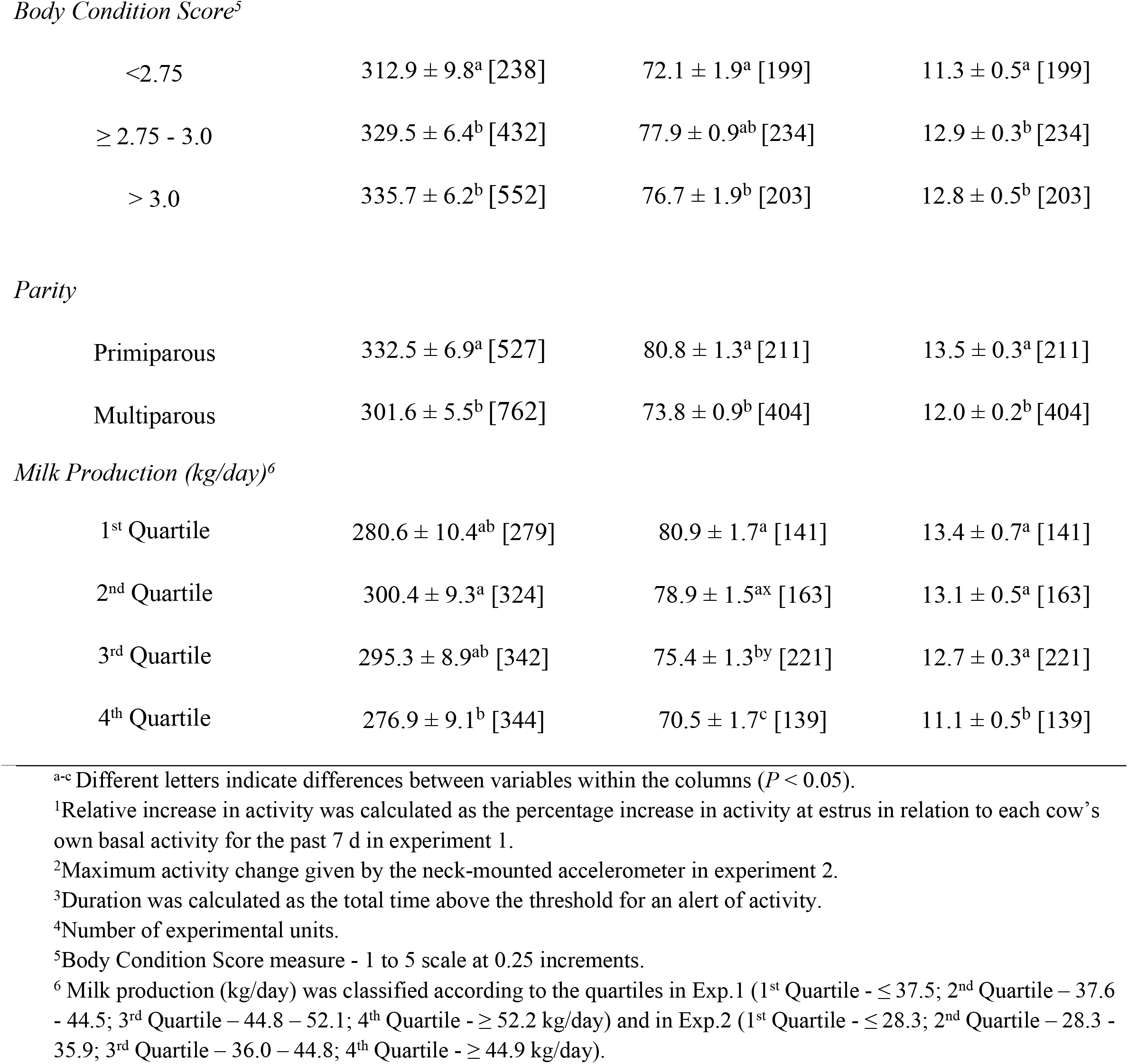
Least squares means ± SEM for relative increase in activity (%), maximum activity change (index) and total duration (h) on body condition score, parity and milk production, in both Experiment 1 and Experiment 2.

| Variables | Relative Increase<sup>1</sup> (%)<br>[n]<sup>4</sup> | Maximum Activity<sup>2</sup><br>Change (index) [n] | Total Duration<sup>3</sup> (h)<br>[n] |
| --- | --- | --- | --- |
| *Body Condition Score*<sup>5</sup> |  |  |  |
| <2.75 | 312.9 ± 9.8<sup>a</sup> [238] | 72.1 ± 1.9<sup>a</sup> [199] | 11.3 ± 0.5<sup>a</sup> [199] |
| ≥ 2.75 - 3.0 | 329.5 ± 6.4<sup>b</sup> [432] | 77.9 ± 0.9<sup>ab</sup> [234] | 12.9 ± 0.3<sup>b</sup> [234] |
| > 3.0 | 335.7 ± 6.2<sup>b</sup> [552] | 76.7 ± 1.9<sup>b</sup> [203] | 12.8 ± 0.5<sup>b</sup> [203] |
| *Parity* |  |  |  |
| Primiparous | 332.5 ± 6.9<sup>a</sup> [527] | 80.8 ± 1.3<sup>a</sup> [211] | 13.5 ± 0.3<sup>a</sup> [211] |
| Multiparous | 301.6 ± 5.5<sup>b</sup> [762] | 73.8 ± 0.9<sup>b</sup> [404] | 12.0 ± 0.2<sup>b</sup> [404] |
| *Milk Production (kg/day)*<sup>6</sup> |  |  |  |
| 1<sup>st</sup> Quartile | 280.6 ± 10.4<sup>ab</sup> [279] | 80.9 ± 1.7<sup>a</sup> [141] | 13.4 ± 0.7<sup>a</sup> [141] |
| 2<sup>nd</sup> Quartile | 300.4 ± 9.3<sup>a</sup> [324] | 78.9 ± 1.5<sup>ax</sup> [163] | 13.1 ± 0.5<sup>a</sup> [163] |
| 3<sup>rd</sup> Quartile | 295.3 ± 8.9<sup>ab</sup> [342] | 75.4 ± 1.3<sup>by</sup> [221] | 12.7 ± 0.3<sup>a</sup> [221] |
| 4<sup>th</sup> Quartile | 276.9 ± 9.1<sup>b</sup> [344] | 70.5 ± 1.7<sup>c</sup> [139] | 11.1 ± 0.5<sup>b</sup> [139] |

<sup>a-c</sup> Different letters indicate differences between variables within the columns ( $P < 0.05$ ).
<sup>1</sup>Relative increase in activity was calculated as the percentage increase in activity at estrus in relation to each cow's own basal activity for the past 7 d in experiment 1.
<sup>2</sup>Maximum activity change given by the neck-mounted accelerometer in experiment 2.
<sup>3</sup>Duration was calculated as the total time above the threshold for an alert of activity.
<sup>4</sup>Number of experimental units.
<sup>5</sup>Body Condition Score measure - 1 to 5 scale at 0.25 increments.
<sup>6</sup> Milk production (kg/day) was classified according to the quartiles in Exp.1 (1<sup>st</sup> Quartile - ≤ 37.5; 2<sup>nd</sup> Quartile - 37.6 - 44.5; 3<sup>rd</sup> Quartile - 44.8 - 52.1; 4<sup>th</sup> Quartile - ≥ 52.2 kg/day) and in Exp.2 (1<sup>st</sup> Quartile - ≤ 28.3; 2<sup>nd</sup> Quartile - 28.3 - 35.9; 3<sup>rd</sup> Quartile - 36.0 - 44.8; 4<sup>th</sup> Quartile - ≥ 44.9 kg/day).

### Pre-ovulatory follicle diameter

Mean pre-ovulatory follicle diameter was 13.4 ± 3.3 mm and 18.7 ± 4.7 mm for Exp. 1 and Exp. 2, respectively. There was no correlation between follicle diameter and relative increase in activity in Exp. 1 (r = -0.01; *P* = 0.89) or maximum activity index (r = - 0.04; *P* = 0.31) and duration (r = - 0.03; *P* = 0.52) in Exp. 2.

In Exp. 1, cows that had higher concentrations of P4 at –4 d had larger pre-ovulatory follicles (14.8 ± 0.2 vs. 12.3 ± 0.3 mm; *P* < 0.001). There was no association of concentration of P4 at AI and the size of the pre-ovulatory follicle size within either study (Exp. 1: 12.9 ± 0.7 vs. 13.0 ± 0.6; *P* = 0.91; Exp. 2.: 17.7 ± 0.5 vs. 18.6 ± 0.5; *P* = 0.20). Cows with higher concentrations of P4 at 7 d post-AI had larger pre-ovulatory follicles at the time of AI within both experiments (Exp. 1: 13.7 ± 0.3 vs. 12.6 ± 0.3 mm; *P* < 0.001; Exp. 2: 19.1 ± 0.5 vs. 17.2 ± 0.5 mm; *P* < 0.001). However, no associations were found between concentrations of P4 at 14 d (*P* = 0.88) or 21 d (*P* = 0.73) post-AI and pre-ovulatory follicle diameter. In Exp. 2, cows that had greater concentrations of E2 at estrus alert had larger pre-ovulatory follicles compared with cows that had lower concentrations of E2 (19.5 ± 0.4 vs. 18.2 ± 0.3 mm; *P* < 0.001).

### Concentration of progesterone and estradiol

Concentrations of P4 at -4 d, 0 d and 7 d post-AI were all associated with the intensity of estrous expression within the current study. In Exp. 1, cows with higher concentrations of P4 at -4 d had greater estrous expression compared with cows that had lower concentrations (318.2 ± 10.5 vs. 289.4 ± 10.6 % relative increase; *P* = 0.05). Lower concentrations of P4 on 0 d were associated with increased intensity and duration of estrous expression (Exp. 1-363.6 ± 5.2 vs. 275.9 ± 8.0 % relative increase; *P* < 0.001; Exp 2 - 76.7 ± 1.9 vs. 67.4 ± 4.7 index; *P* < 0.05; Exp. 2 - 12.5 ± 0.5 vs. 9.3 ± 1.8 h; *P* < 0.001). Similarly, cows that had lower intensity and shorter duration of estrous expression not only had higher concentrations of P4 but also lower concentrations of E2 at estrus compared with cows with greater activity and longer duration estrous behaviour (Figure 2A and Figure 2B; *P* < 0.01). The proportion of cows displaying greater and lower estrous intensity was shifted between cows with higher or lower concentrations of P4 at -4 d in Exp. 1 (Figure 3A) and 0 d in both, Exp. 1 and Exp. 2 (Figure 3B). Moreover, a total of 66.7% of the cows with greater concentration of P4 at -4 d had lower concentration of P4 at 0 d. Together, cows classified as HL had greater estrous intensity when compared with cows classified as LH, HH and LL (356.6 ± 5.5 vs. 272.4 ± 7.4 vs. 293.8 ± 4.8 vs. 267.2 ± 14.6 % relative increase; *P* < 0.001), whereas the latter three categories did not differ among groups.

**Figure 2.**
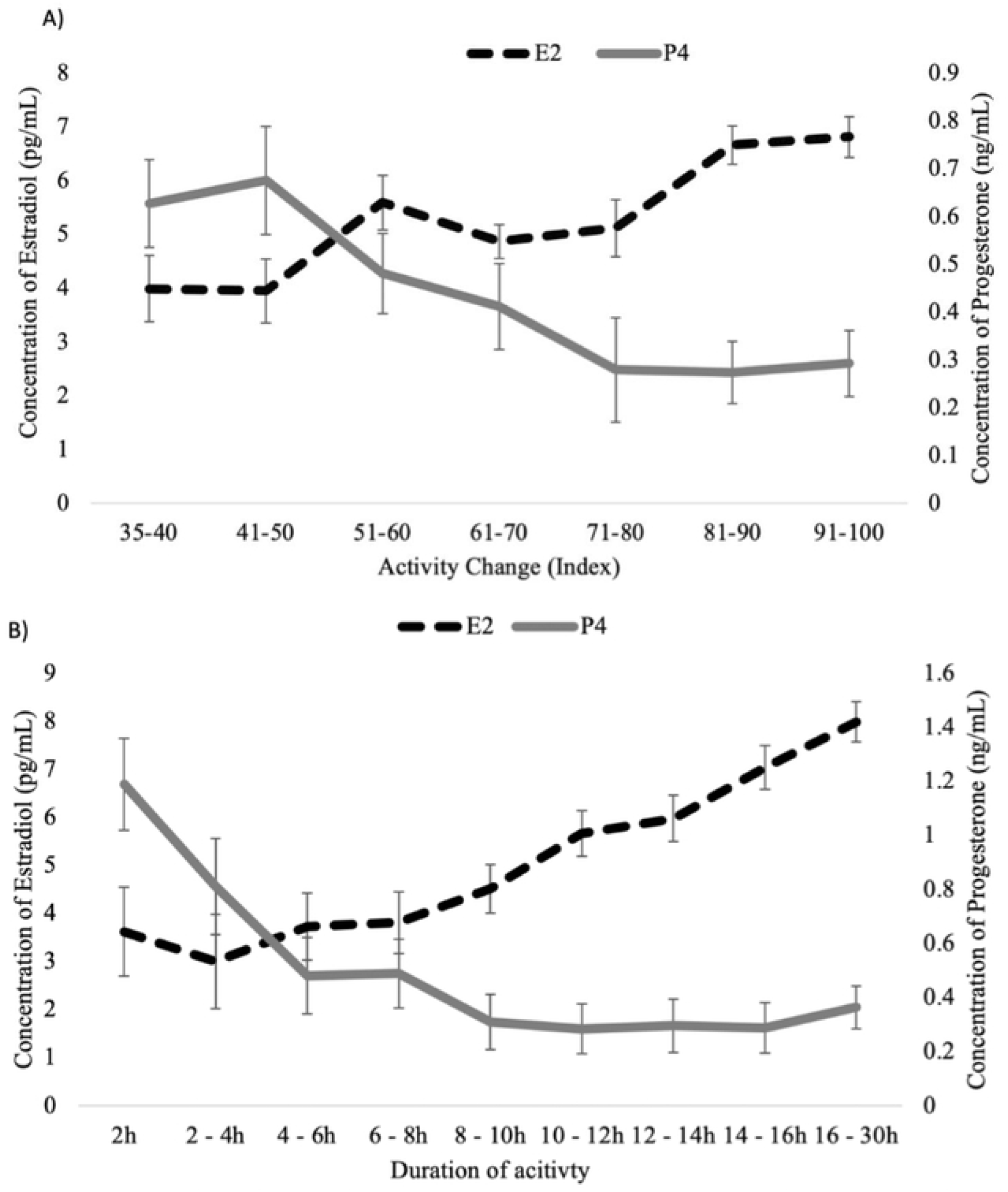
Distribution of increased estrous expression intensity (panel A) and duration (panel B) detected by a neck-mounted accelerometer at estrus and its association with the concentrations of progesterone and estradiol (least squares 1neans ± SE) in Experi1nent 2.

**Figure 3.**
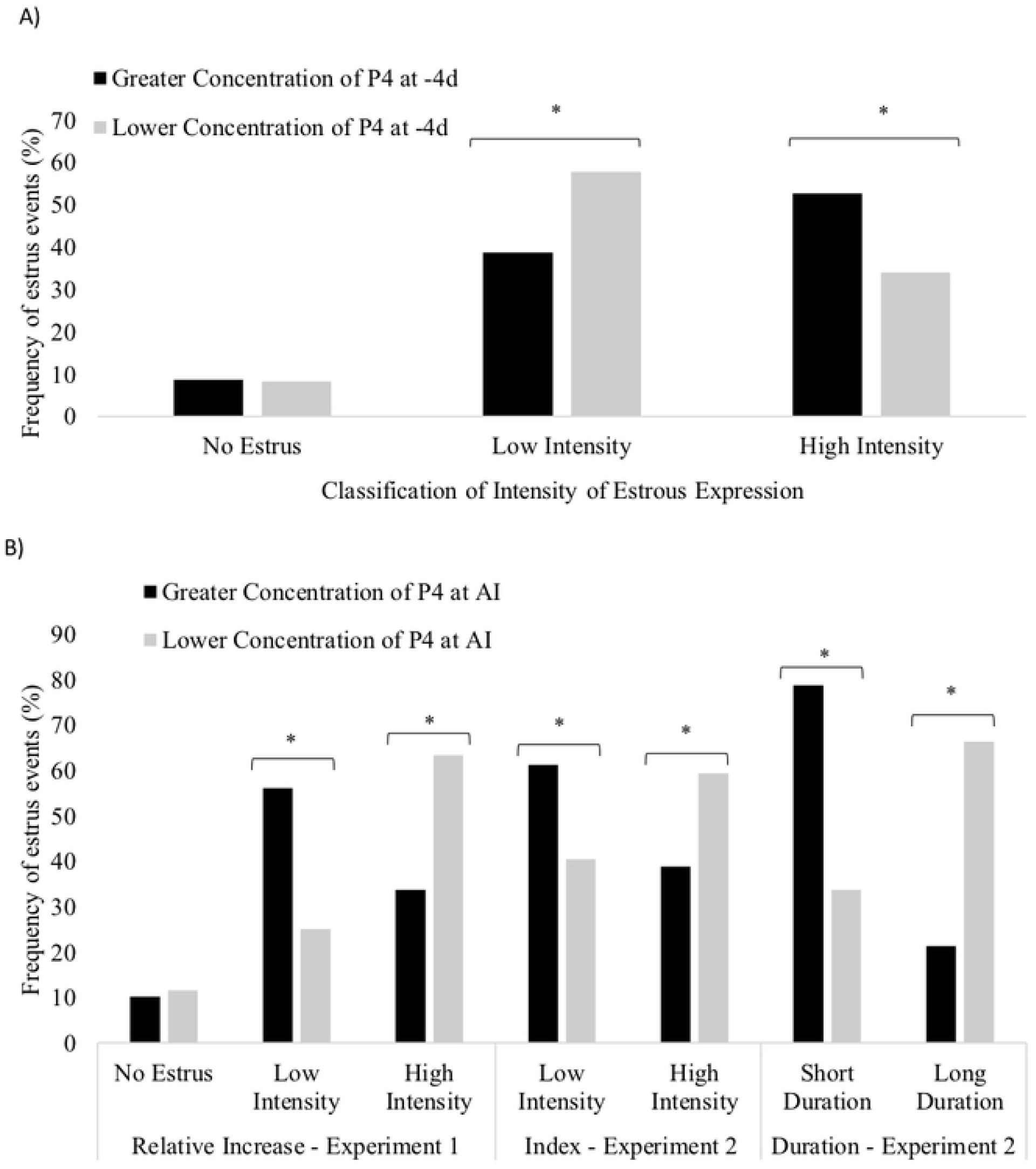
Proportion of cows displaying greater and lesser estrous expression according to the concentration of P4 at -4 d (Experiment I, Panel A; *P* < 0.01), and the concentration of P4 at Od (Experiment I and Experi1nen1 2, Panel B; *P* < 0.0 I).

Cows with higher intensity of estrus at timed AI had greater concentrations of P4 at 7 d post-AI compared with those that had lower intensity or those that did not express estrus (3.45 ± 0.1 vs. 3.19 ± 0.1 vs. 2.16 ± 0.2 ng/mL; *P* < 0.001). Greater concentrations of P4 on 7 d (*P* < 0.05), 14 d (*P* < 0.01) and 21 d (*P* < 0.01) post-AI were found in cows that had higher intensity of estrus detected at the time of AI (Figure 3). Size of the corpus luteum on 7 d (*P* = 0.83), 14 d (*P* = 0.67) and at 21 d (*P* = 0.58) post-AI did not differ between cows that expressed higher or lower intensity of estrus.

Milk production was not correlated with concentrations of P4 at -4 d (r = 0.01; P = 0.75), 0 d (Exp. 1: r = 0.02; *P* = 0.51; Exp. 2: r = - 0.01; P = 0.91), 7 d (Exp. 1: r = 0.05; *P* = 0.12; Exp. 2: r = - 0.05; *P* = 0.50), 14 d (r = 0.006; *P* = 0.94) and 21 d post-AI (r = 0.004; *P* = 0.95) or with the concentration of E2 on 0 d (r = - 0.09; P = 0.31). Concentrations of P4 at AI and 7 d post-AI, relative to BCS, parity and milk production, are summarized in Table 2 for both Exp. 1 and 2.

**Table 2.**
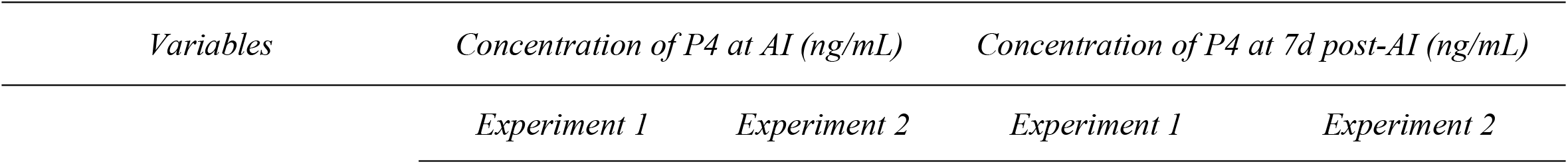
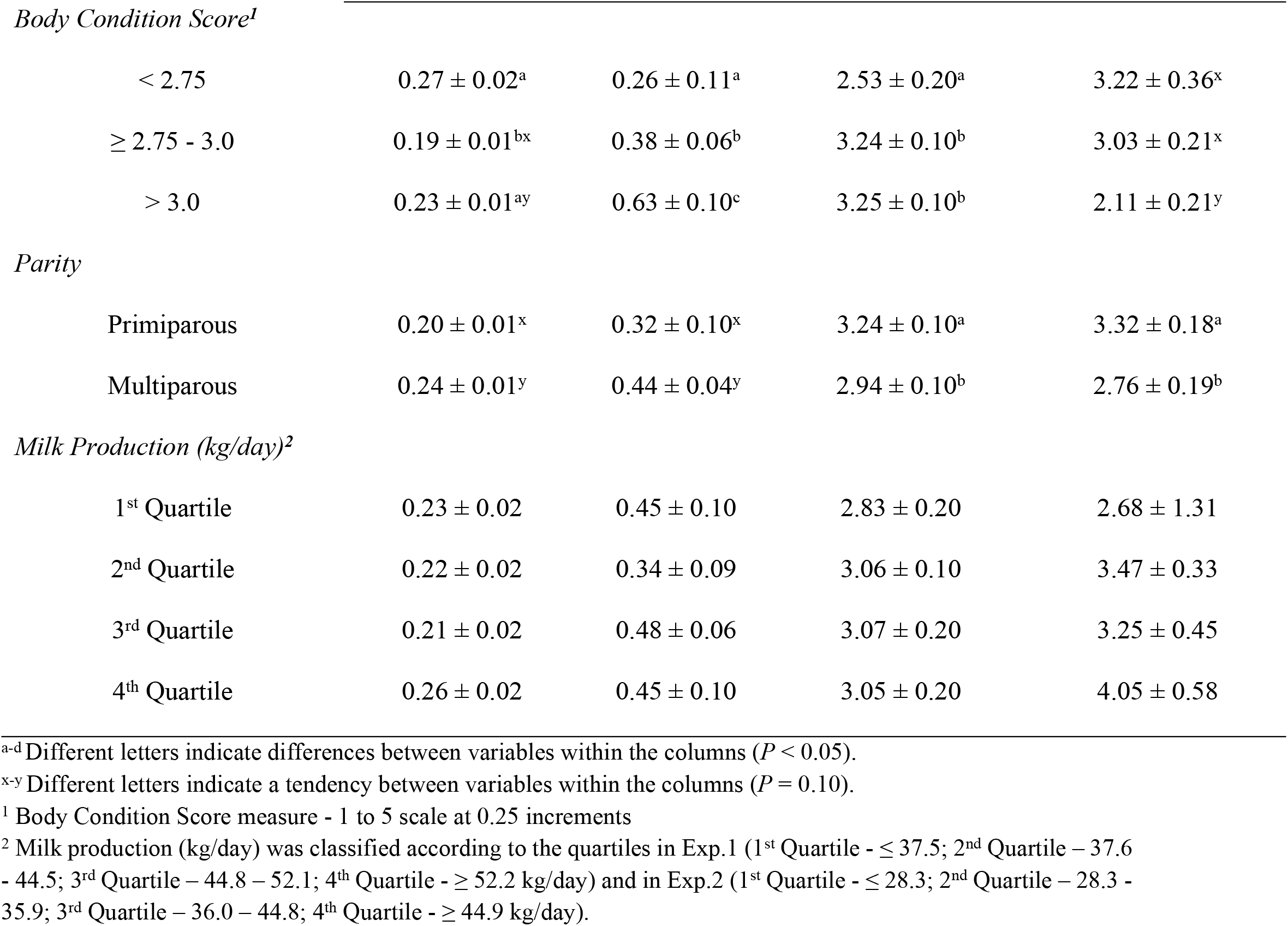
Concentration of progesterone (least squares means ± SE) at AI and at 7 d post-AI, according to body condition score, parity and milk production on both Experiment 1 and Experiment 2.

| *Variables* | *Concentration of P4 at AI (ng/mL)* |  | *Concentration of P4 at 7d post-AI (ng/mL)* |  |
| --- | --- | --- | --- | --- |
|  | *Experiment 1* | *Experiment 2* | *Experiment 1* | *Experiment 2* |
| *Body Condition Score<sup>1</sup>* |  |  |  |  |
| < 2.75 | 0.27 ± 0.02<sup>a</sup> | 0.26 ± 0.11<sup>a</sup> | 2.53 ± 0.20<sup>a</sup> | 3.22 ± 0.36<sup>x</sup> |
| ≥ 2.75 - 3.0 | 0.19 ± 0.01<sup>bx</sup> | 0.38 ± 0.06<sup>b</sup> | 3.24 ± 0.10<sup>b</sup> | 3.03 ± 0.21<sup>x</sup> |
| > 3.0 | 0.23 ± 0.01<sup>ay</sup> | 0.63 ± 0.10<sup>c</sup> | 3.25 ± 0.10<sup>b</sup> | 2.11 ± 0.21<sup>y</sup> |
| *Parity* |  |  |  |  |
| Primiparous | 0.20 ± 0.01<sup>x</sup> | 0.32 ± 0.10<sup>x</sup> | 3.24 ± 0.10<sup>a</sup> | 3.32 ± 0.18<sup>a</sup> |
| Multiparous | 0.24 ± 0.01<sup>y</sup> | 0.44 ± 0.04<sup>y</sup> | 2.94 ± 0.10<sup>b</sup> | 2.76 ± 0.19<sup>b</sup> |
| *Milk Production (kg/day)<sup>2</sup>* |  |  |  |  |
| 1<sup>st</sup> Quartile | 0.23 ± 0.02 | 0.45 ± 0.10 | 2.83 ± 0.20 | 2.68 ± 1.31 |
| 2<sup>nd</sup> Quartile | 0.22 ± 0.02 | 0.34 ± 0.09 | 3.06 ± 0.10 | 3.47 ± 0.33 |
| 3<sup>rd</sup> Quartile | 0.21 ± 0.02 | 0.48 ± 0.06 | 3.07 ± 0.20 | 3.25 ± 0.45 |
| 4<sup>th</sup> Quartile | 0.26 ± 0.02 | 0.45 ± 0.10 | 3.05 ± 0.20 | 4.05 ± 0.58 |

<sup>a-d</sup> Different letters indicate differences between variables within the columns ( $P < 0.05$ ).
<sup>x-y</sup> Different letters indicate a tendency between variables within the columns ( $P = 0.10$ ).
<sup>1</sup> Body Condition Score measure - 1 to 5 scale at 0.25 increments
<sup>2</sup> Milk production (kg/day) was classified according to the quartiles in Exp.1 (1<sup>st</sup> Quartile - ≤ 37.5; 2<sup>nd</sup> Quartile - 37.6 - 44.5; 3<sup>rd</sup> Quartile - 44.8 - 52.1; 4<sup>th</sup> Quartile - ≥ 52.2 kg/day) and in Exp.2 (1<sup>st</sup> Quartile - ≤ 28.3; 2<sup>nd</sup> Quartile - 28.3 - 35.9; 3<sup>rd</sup> Quartile - 36.0 - 44.8; 4<sup>th</sup> Quartile - ≥ 44.9 kg/day).

### Pregnancy per AI (P/AI)

In Exp. 1, P/AI was influenced by estrous expression, parity, BCS, concentration of P4 at - 4 d, at 0 d and 7 d post-AI, but not by milk production. Cows that expressed greater intensity of estrus had greater P/AI compared with those with lower intensity or those that did not express estrus at timed AI (46.2 ± 3.0 % vs. 33.6 ± 3.6 % vs. 5.4 ± 8.1 %; *P* < 0.01). The concentration of P4 at -4 d, at the day of timed AI (0 d) and 7 d post-AI and its association with P/AI is shown in Table 3. Altogether, cows classified as HL had higher P/AI when compared with cows classified as LH, LL and HH (31.2 ± 2.1 % vs. 18.1 ± 2.8 % vs. 20.5 ± 4.1 % vs. 9.7 ± 4.9 %; *P* < 0.001). Parity and BCS were associated with P/AI. Multiparous cows had reduced P/AI compared with primiparous cows (32.2 ± 3.6 % vs. 46.5 ± 4.1 %; *P* < 0.01) and cows with high and moderate BCS had greater P/AI than cows with Low BCS (36.1 ± 1.9 % vs. 31.7 ± 2.0 % vs. 22.1 ± 3.0 %; *P* < 0.001). No interaction between estrus intensity and parity was found on P/AI (*P* = 0.37); however, there was an interaction between estrus intensity and BCS on P/AI (*P* = 0.05). Milk production had no effect on P/AI (*P* = 0.44).

**Table 3.** Pregnancy per AI, in Experiment 1, according to the concentration of progesterone at – 4 d, at the day of AI (0 d) and 7 d post timed AI.

| Variables | Pregnancy per AI (%) [n/n] |
| --- | --- |
| *Concentration of P4 at – 4 d\** |  |
| $\leq 3.04$ ng/mL | $22.4 \pm 4.5^a$ [34/151] |
| > 3.04 ng/mL | 32.8 ± 4.4<sup>b</sup> [53/153] |
| *Concentration of P4 at AI (0 d) \** |  |
| ≤ 0.1 ng/mL | 35.2 ± 3.4<sup>a</sup> [214/467] |
| > 0.1 ng/mL | 19.6 ± 4.6<sup>b</sup> [40/143] |
| *Concentration of P4 at 7 d post-AI \** |  |
| ≤ 2.96 ng/mL | 39.1 ± 2.9<sup>a</sup> [169/390] |
| > 2.96 ng/mL | 24.7 ± 2.6<sup>b</sup> [108/387] |

<sup>a-b</sup> Different letters indicate differences between variables within the columns ( $P < 0.05$ ).
\*Concentrations of P4 were classified as lesser or equal to the median and greater than the median at each sampling time using the following values: - 4 d prior to timed AI (3.04 ng/mL); 0 d (at the moment of timed AI; 0.1 ng/mL) and at 7 d post-AI (2.96 ng/mL).

In Exp. 2, P/AI was influenced by estrous expression, BCS, and concentration of P4 at 0d and tended to be influenced by concentrations of P4 at 7 d, 14 d and 21d post-AI, but not by milk production or parity. Cows that expressed higher intensity or longer duration estrous expression had greater P/AI compared to those with lower intensity or shorter duration estrous expression at AI (High intensity: 39.9 ± 3.6 % vs. Low intensity: 26.8 ± 2.6 %; *P* < 0.001; Long duration: 31.2 ± 2.5 % vs. Short duration: 23.1 ± 3.2 %; *P* < 0.001). Cows with lesser concentrations of P4 at 0 d (≤ 0.26 ng/mL) had greater P/AI compared with cows with greater concentrations (31.8 ± 2.8 % vs. 23.4 ± 3.2 %; *P* < 0.01). Cows that had lower concentrations of P4 at 7 d, 14 d and 21 d post-AI tended to have lower P/AI when compared with cows with greater concentrations of P4 (*P* = 0.10; Figure 4). Parity was not associated with pregnancy (*P* = 0.83) in Exp. 2. Body condition score was associated with P/AI (*P* < 0.001), as cows with High and Moderate BCS had greater P/AI than cows with Low BCS (32.9 ± 2.9 % vs. 42.3 ± 4.6 % vs. 19.5 ± 4.7 %; *P* < 0.001). An interaction between estrus intensity and parity on P/AI was found (*P* < 0.001), as well as a tendency for an interaction between estrus intensity and BCS on P/AI (*P* = 0.10), as shown in Table 4. Milk production had no association with P/AI (*P* = 0.80).

**Table 4.**
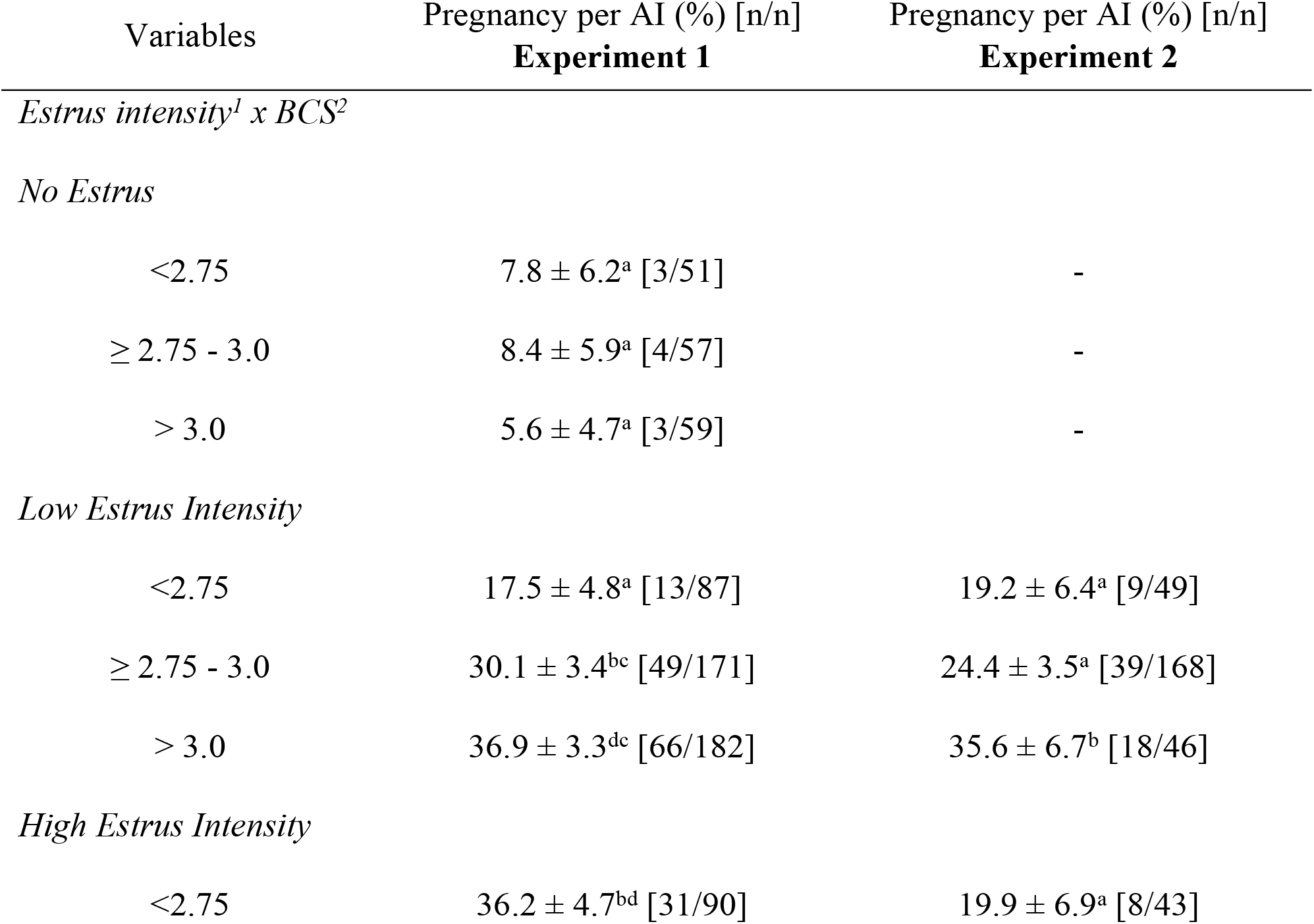
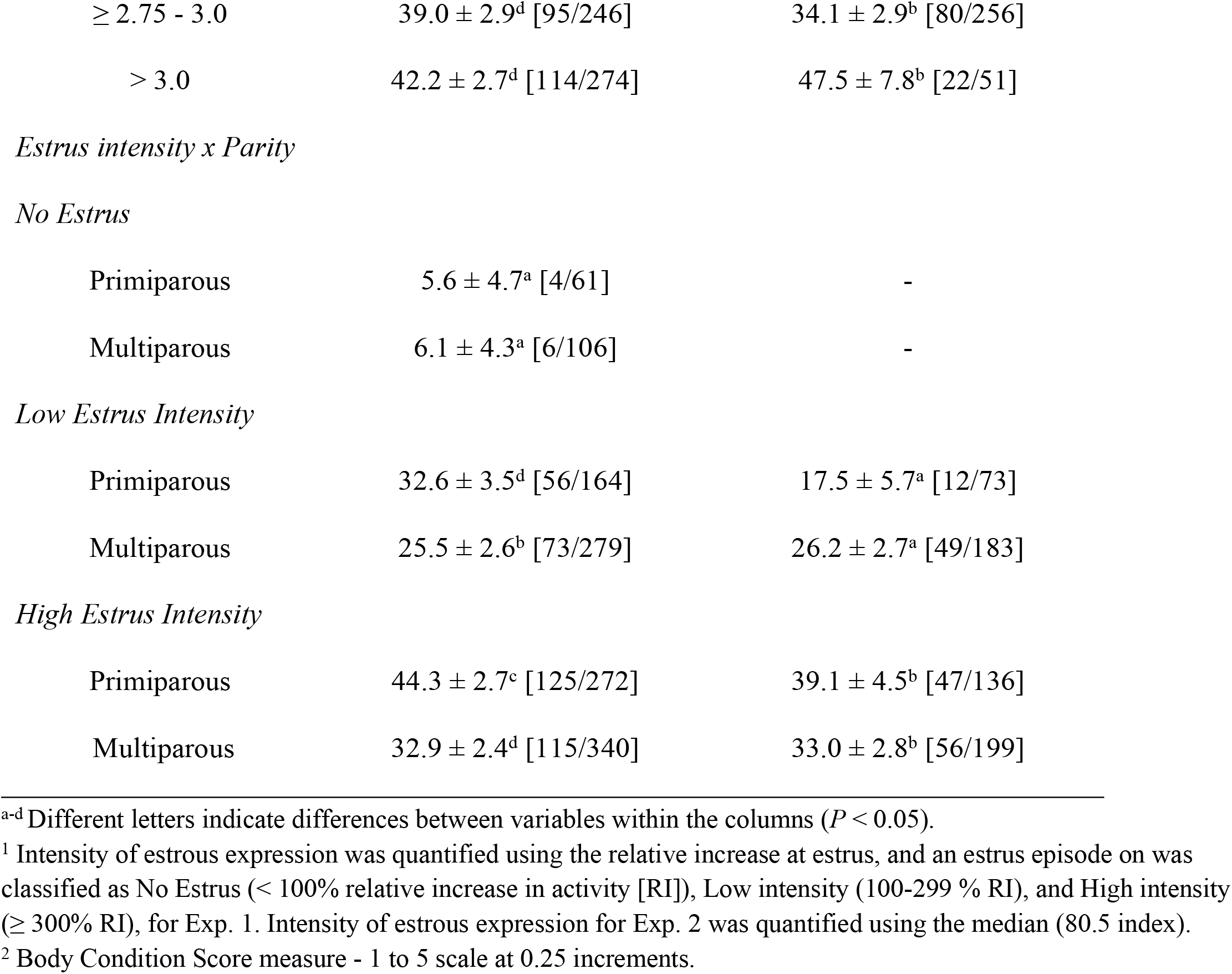
Pregnancy per AI, in both Experiment 1 and Experiment 2, according to the interactions of estrus intensity and body condition score and the interaction of estrus intensity and parity.

**Figure 4.**
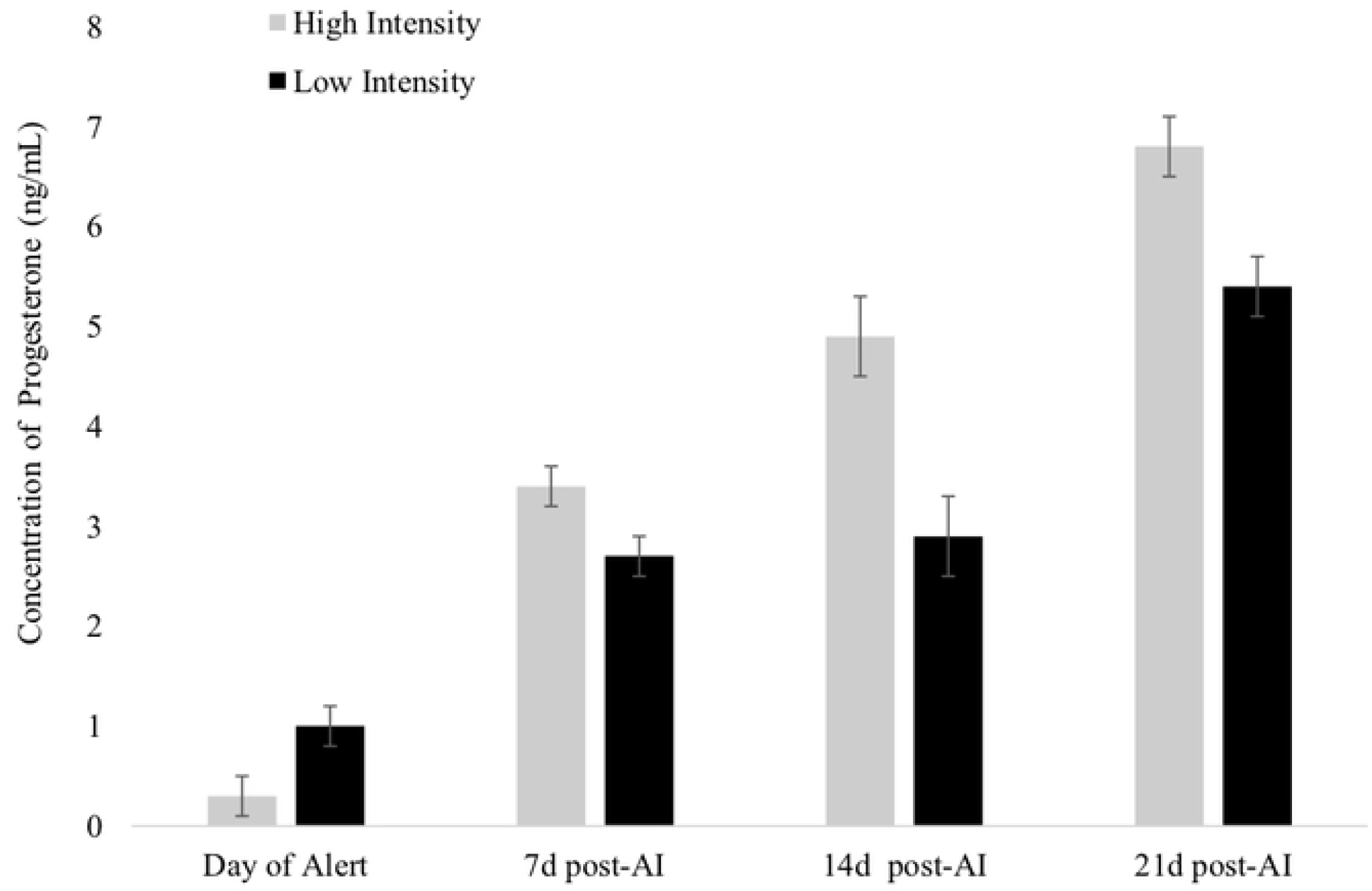
Concentrations of progesterone (least squares means± SE) on the day of estrus alert (0 d; *P* < 0.01) and 7 d (*P* <0.0S), 14 d (*P* <0.01), and 21 d (*P* <0.01) post-Al relative to estrous expression intensity in Experiment 2. High intensity: estrous expression greater or equal to the median (80.5 index) of the automated activity monitor. Low intensity: estrous expression less than the median.

**Figure 5.**
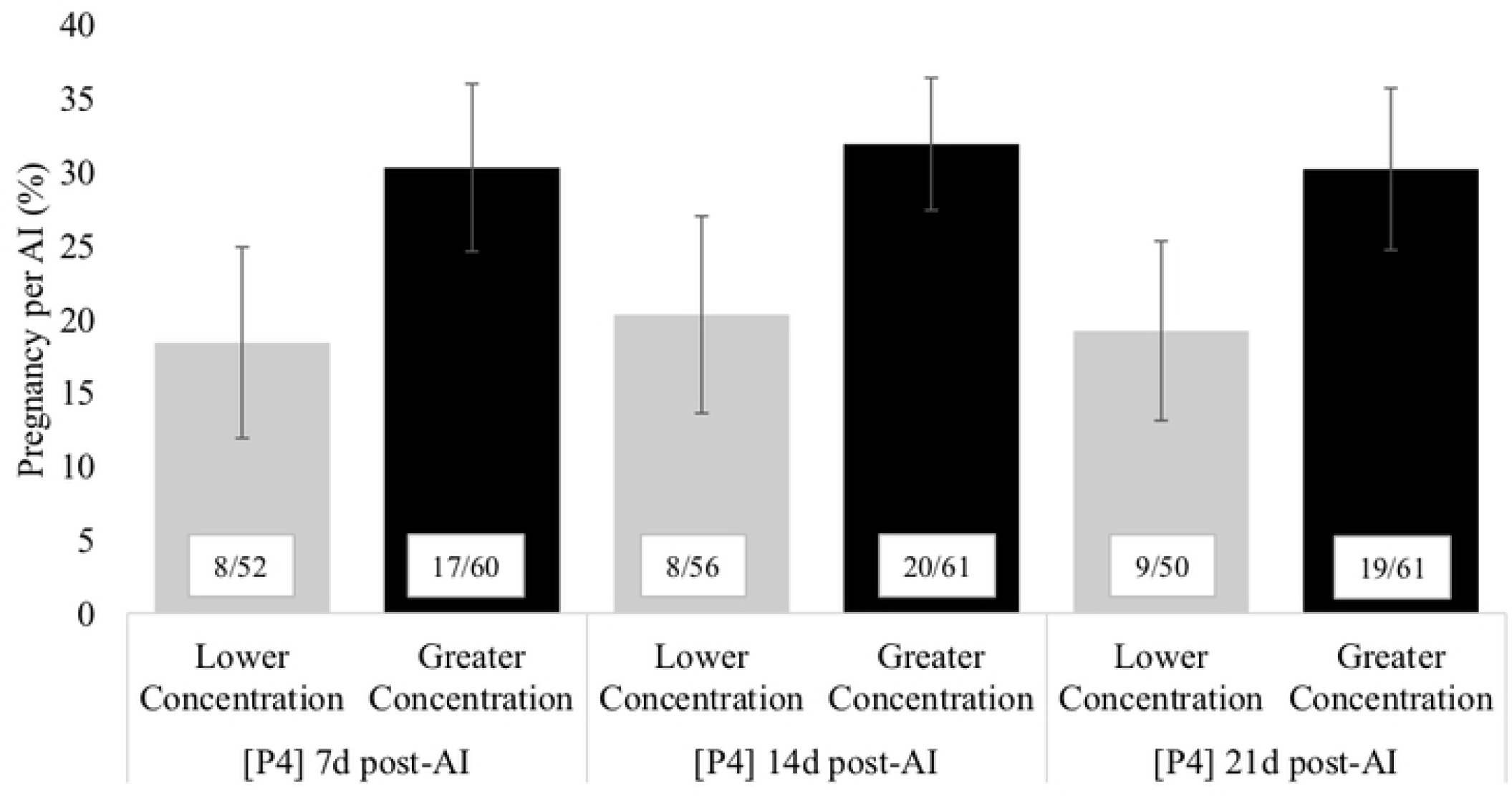
Pregnancy per artificial insemination (AI;%) according to the concentration of progesterone (P4) on days 7,14 and 21 d post-Al (*P* = 0.10). Concentration of P4 was classified as higher and lower concentration according to the median as follow: 7 d post-Al= ≤ 2.65 and > 2.65 ng/rnL; 14 d post-Al= ≤ 3.12 and > 3.12 ng/mL; and 21 d post-Al= ≤ 6.27ng/mL and > 6.27ng/mL).

## Discussion

The overall objective of this study was to evaluate whether the concentration of P4 at different phases of the estrous cycle was associated the intensity of estrous expression considering both spontaneous estrus and a timed AI protocol and to assess the association with P/AI. Overall, we found that cows with more ideal hormonal concentrations had higher intensity of estrous expression and greater P/AI. Cows with greater concentrations of P4 just before luteolysis and on 7d post-AI were more likely to have had greater intensity estrous expression at AI. Greater concentrations of P4 (Exp. 1 and 2) and lower concentrations of E2 (Exp. 2) at AI were associated with lower intensity estrous expression, detected by AAM. Altogether, the findings from the current study suggest that greater concentrations of P4 during the growth of the preovulatory follicle, and a greater concentration of P4 during the early stages of embryo development are closely associated with the intensity of induced and spontaneous estrous expression. In addition, lower concentrations of P4 at AI, even in induced estrus events, was linked with estrous expression and fertility, suggesting an association between complete CL regression and the behavioural trigger of estrus in the hypothalamus.

Greater concentrations of P4 at -4 d were associated with a greater intensity of estrous expression and increased fertility. It has been shown that P4 primes the hypothalamus to be more responsive to E2 [23], due to increased expression of estradiol receptors in the hypothalamus [24,25]. Walker et al. [26] reported that low P4 exposure before estrus was associated with low intensity of sexual behaviors during estrus in chronically stressed lame cows. Progesterone may cause up- or downregulation in the hypothalamus of a number of genes that are involved in the estrous behavior through E2 receptors [27]. A study by Herlihy et al. [28] demonstrated that lower concentrations of P4 during diestrus was associated with lower fertility. Progesterone can block the estrus-inducing actions of E2 and plays an important role in priming the bovine brain for E2 functions [29,30]. Increased estrous expression in timed AI protocols that include supplementation of P4 also suggests that P4 may act as a primer for the responsiveness of the hypothalamus to E2 [31]. Bisinotto et al. [32] compared different timed AI protocols and reported that an addition of an intravaginal progesterone device in the first wave of follicle development increased the proportion of cows inseminated in estrus at the end of the timed AI.

In both experiments greater concentrations of P4 at AI were associated with lower intensity and duration of estrus and lower fertility. Pereira et al. [8] reported that cows that did not express estrus at the end of a timed AI protocol were more likely to have higher concentrations of P4 compared with cows that expressed estrus. Higher concentrations of P4 at the time of AI have also been associated with lower fertility [8]. A physiological mechanism that may reduce fertility when P4 is elevated at AI is that P4 may alter oocyte transport by altering uterine or oviductal contractility, thus reducing fertilization [33]. A study in rats demonstrated that the facilitation of sperm migration into the oviduct is negatively affected by P4 [34]. Cerri et al. [35] suggested that cows with low P4 prior to AI had increased basal LH concentrations, altering follicular dynamics that could affect oocyte quality. Another hypothesis to explain lower fertility is that P4 could have detrimental impacts on embryo quality and development. When P4 was added to an *in vitro* fertilization protocol, it reduced the blastocyst percentage, suggesting that P4 may have a direct effect on early embryonic development [36]. In addition, slight elevations in P4 are associated with reduced endometrial thickness [36], suggesting that the effects of P4 on the uterus could also reduce embryo development.

In both experiments, cows with a higher increase in estrous expression had higher concentrations of P4 post-AI (Exp.1 – 7 d post-AI and Exp.2 – 7 d, 14 d and 21 d post-AI) and greater P/AI. The concentration of P4 post-AI has been shown to affect fertility in lactating dairy cows [37–39] and is a requirement for establishment of pregnancy [40]. This positive effect of P4 post-AI on fertility has been associated with the elongation of the conceptus [41], early embryonic development [42], and increased secretion of interferon-tau [43]. Mann et al. [44] observed that the insertion of an intravaginal P4 device between day 5 and 9 of the estrous cycle caused an increase in embryo length 16 days after AI.

The diameter of the pre-ovulatory follicle during the estrous cycle may also be a factor that contributes to concentrations of P4 after AI, as follicles with larger diameters have been shown to generate larger CLs, resulting in higher endogenous production of P4 [45]. However, other studies have not always observed this relationship, as some have reported no difference in the concentration of P4 post-ovulation, even with differences in follicular diameter and the formation of CLs with larger diameters [35,46].

One important finding of the current study is that the combination of P4 concentrations at different times of the estrous cycle is likely more important than just one measurement alone. For example, it was observed that combining different categories of -4 d and 0 d, P4 concentrations can be associated with estrous expression and fertility outcomes. There is still need for more randomly controlled trials to properly suggest cause-effect, but there is increasing evidence of the effect of P4 on the trigger and intensity of estrus. One study [47] using long or short exposure times to P4 during the follicular growth phase, demonstrated that longer exposure of P4 was associated with numerically higher estrous expression. Animal related factors such as parity, BCS, and milk production have been consistently associated with estrous expression. In all cases, primiparous cows with higher BCS were more likely to express high intensity estrus; these cows were also more likely to have greater concentrations of circulating P4 and improve fertility [48].

## Conclusion

This research demonstrates that P4 concentrations around the time of AI are important for the expression of estrus for both timed AI and AI based on spontaneous estrus. As the intensity of estrus is associated with greater P/AI, future studies should determine if cows can be selected for their ability to maintain hormonal profile (i.e., high concentration of P4 prior to AI, low concentration of P4 at AI and high concentration of P4 post-AI) associated with increased estrous expression and fertility.

## Acknowledgments

This study was supported by a contribution from the Dairy Research Cluster Initiative II (Dairy Farmers of Canada, Agriculture and Agri-Food Canada, the Canadian Dairy Network and the Canadian Dairy Commission; Ottawa, Canada). Partial funding was also provided by the Natural Sciences and Engineering Research Council, Discovery Grant (Ottawa, Canada). This study was a part of the Resilient Dairy Genome Project (RDGP). We gratefully acknowledge the funding and support from the Resilient Dairy Genome Project including Genome Canada, Genome Alberta, Genome BC. We are thankful to Fazenda Colorado (Araras, Brazil) and the UBC Dairy Education and Research Centre (Agassiz, Canada) and their farm personnel for contributing with this research project as well as students, interns and operational support from Conapec Jr., Sao Paulo State University, Botucatu, Brazil.

